# Structural and functional characterization of ubiquitin variant inhibitors for the JAMM-family deubiquitinases STAMBP and STAMBPL1

**DOI:** 10.1101/2021.07.16.452720

**Authors:** Yusong Guo, Qi Liu, Evan Mallette, Cody Caba, Feng Hou, Julia Fux, Gabriel LaPlante, Aiping Dong, Qi Zhang, Hui Zheng, Yufeng Tong, Wei Zhang

**Author notes:** These authors contributed equally to this work (Yusong Guo, Qi Liu, Evan Mallette). Corresponding authors: (YT) and (WZ).

## Abstract

Ubiquitination is one of the most crucial post-translational protein modifications involved in a myriad of biological pathways. This process is reversed by deubiquitinases (DUBs) that deconjugate the ubiquitin (Ub) moiety or poly-Ub chains from substrates. In the past decade, tremendous efforts have been focused on targeting DUBs for drug discovery. However, most chemical compounds with inhibitory activity for DUBs suffer from mild potency and low selectivity. To overcome these obstacles, we developed a phage display-based protein engineering strategy for generating Ub variant (UbV) inhibitors and have previously successfully applied it to the Ub-specific protease (USP) family of cysteine proteases. In this work, we leveraged the UbV platform to target STAMBP, a member of the JAB1/MPN/MOV34 (JAMM) metalloprotease family of DUB enzymes. We identified two UbVs (UbV^SP.1^ and UbV^SP.3^) that bind to STAMBP with high affinity but differ in their selectivity for the closely related paralog STAMBPL1. We determined the STAMBPL1-UbV^SP.1^ complex structure by X-ray crystallography, revealing hotspots of the tight JAMM-UbV interaction. Finally, we show that UbV^SP.1^ and UbV^SP.3^ are potent inhibitors of the STAMBP isopeptidase activity, far exceeding the reported small-molecule inhibitor BC-1471. This work demonstrates that UbV technology is suitable to develop tool molecules for metalloproteases. These tools can be used to understand the cellular function of JAMM family DUBs.

## Introduction

Ubiquitin (Ub) is a small, highly conserved protein that plays a central role in the Ub-proteasome system (UPS) to tightly regulate numerous biological processes, including immune responses (1), DNA repair (2), and the cell cycle (3). During ubiquitination, the C-terminal carboxyl group of Ub is covalently attached to a substrate protein at the lysine ε-amine or the N-terminal primary amine through an isopeptide or an α-peptide bond, respectively. The conjugated Ub itself can be further ubiquitinated at its N-terminus or one of seven lysine residues, forming a poly-Ub chain (4). The ubiquitination process is catalyzed by a three-enzyme cascade comprised of E1 activating enzymes, E2 conjugating enzymes, and E3 Ub ligases. Different linkages of poly-Ub chains serve distinct purposes. For example, the most abundant K48-linked poly-Ub chains target substrate proteins to proteasomal degradation (4), whereas K63-linked poly-Ub chains are mostly non-degradative and involved in modulating cellular signal transduction (5).

The antagonists of ubiquitination are the deubiquitinase (DUB) enzymes that deconjugate Ub from modified protein substrates. The human genome encodes approximately 100 DUBs, dysregulation of which results in a variety of diseases (6). DUBs are categorized into seven families based on structural folds: ubiquitin-specific proteases (USPs), JAB1/MPN/MOV34 metalloenzymes (JAMMs), ovarian tumor proteases (OTUs), ubiquitin C-terminal hydrolases (UCHs), Machado-Josephin domain-containing proteases (MJDs), motif interacting with ubiquitin-containing novel DUB family (MINDYs) (7, 8), and newly discovered zinc finger containing ubiquitin peptidase (ZUP) (9). DUBs have emerged as attractive targets for pharmacological intervention in various human diseases, particularly cancers (6). However, most small-molecule DUB inhibitors developed have suffered from mild potency and poor selectivity (10). One of the challenges in developing chemical probe quality (11) small-molecule DUB inhibitors is that the Ub-binding groove is usually large, shallow, and not suitably shaped for small-molecule binding (10). The recent success in discovering potent and selective USP7 small-molecule inhibitors took a decade (12–16) and elegantly combined fragment-based screens and structure-guided optimization. This strategy has yet to show general applicability to other members of the DUB superfamily. Here, we set out to develop potent and specific inhibitors for the JAMM-family DUBs.

While the other six families of human DUBs are cysteine proteases, the JAMM family is unique in that they are metalloproteases. The JAMM domain (17) contains a catalytic zinc ion coordinated by two histidines, one aspartate/glutamate, and one water molecule that is hydrogen-bonded to an adjacent glutamate (18). The JAMM family comprises a total of 14 DUBs, seven of which are predicted to be catalytically inactive (pseudo-DUBs) because of substitutions of essential Zn^2+^-coordinating residues (19). Among the seven other JAMM members, STAM-binding protein (STAMBP, also known as AMSH), STAMBP-like 1 (STAMBPL1, also known as AMSH-LP), and MYSM1 (20) can function as isopeptidases independently, whereas BRCA1/BRCA2 containing complex subunit 36 (BRCC36), proteasome 26S subunit, non-ATPase 14 (PSMD14, also known as RPN11), and COP9 signalosome subunit 5 (CSN5) act in macromolecular assemblies (21), and finally, the DUB activity and function of MPND is poorly understood (22).

All known broad-spectrum JAMM-family inhibitors are chelating agents (e.g., 1,10-phenanthroline and thiolutin), which have a high affinity for divalent metal ions and chelate the active site Zn^2+^ ion (23, 24). Notably, a potent and moderately specific RPN11 inhibitor capzimin was developed from a chelating agent-like small molecule through medicinal chemistry optimization (25). CSN5i-3, an orally available inhibitor, has been discovered by Novartis targeting CSN5’s deneddylation activity and confer high specificity towards other metalloproteinases (26). For this work, we chose STAMBP as the representative JAMM DUB for targeted inhibition.

STAMBP consists of three distinct regions, including an N-terminal microtubule interacting and trafficking (MIT) domain, a central SH3-binding motif (SBM), and a C-terminal catalytic JAMM domain (18). STAMBP plays an essential role in endocytosis and endosomal-lysosomal sorting of cell-surface receptors by deubiquitinating and rescuing ubiquitinated cargo proteins from lysosomal degradation. The MIT and SBM domains mediate critical protein-protein interactions (17) to recruit STAMBP to the endosomal sorting complexes required for transport (ESCRT), which regulate the multivesicular body (MVB) biogenesis of ubiquitinated cell-receptors (27). Recessive mutations in the *STAMBP* gene were found to cause a severe developmental disorder, microcephaly capillary malformation syndrome (MIC-CAP), due to elevated Ub-conjugate aggregation and resulted progressive apoptosis (28).

At the biochemical level, STAMBP cleaves K63-linked poly-Ub chains specifically, and the cleavage efficiency is increased when binding to STAM proteins (STAM1 or STAM2) (29). STAMBPL1 shares 56% overall sequence identity to STAMBP and 68% identity in the catalytic domains (30). Both STAMBP and STAMBPL1 localize to early endosomes by binding to clathrin (31); however, STAMBPL1 fails to bind to STAM due to residue substitutions in the SBM-like motif, suggesting it may function in different signaling pathway from that of STAMBP (30). The crystal structure of STAMBPL1^JAMM^ catalytic domain in complex with K63-linked diUb revealed its linkage specificity towards K63-linked poly-Ub chains involves the catalytic groove and two insertions that are conserved in STAMBP subfamily JAMMs: ins-1 (aa 314–339) and ins-2 (aa 393–415) (32). The distal Ub interacts with ins-1 and the catalytic groove, and the proximal Ub interacts with ins-2 and the catalytic site. Recently, Bednash *et al*. used *in silico* molecular modeling and virtual screening and discovered a compound, BC-1471, that inhibits STAMBP and decreases protein level of its inflammasome substrate NALP7. However, the *in vitro* DUB assay showed that BC-1471 could not fully inhibit STAMBP activity even at 100 µM and no cocrystal structure was provided to support the mechanism of action (33).

In this work, we conducted phage-display selections and identified Ub variant (UbV) inhibitors for STAMBP with high binding affinity. We solved the crystal structure of STAMBPL1 in complex with one of two UbVs and identified the hotspots responsible for tight binding. In addition, *in vitro* functional characterization confirmed specific and potent inhibition of STAMBP by UbVs. Importantly, the UbVs exhibited an inhibitory effect superior to BC-1471.

## Results

### Identification of UbV binders for STAMBP

To develop potent and specific inhibitors for STAMBP, we decided to employ a structure-based combinatorial Ub engineering strategy to inhibit the Ub-STAMBP protein-protein interaction (34) selectively. Large hydrophobic surfaces on Ub (~2000 Å^2^) mediate interaction with DUBs, including the Ile36 patch (Ile36, Leu71, and Leu73), the Ile44 patch (Leu8, Ile44, His68, and Val70), and the Phe4 patch (Gln2, Phe4, Thr14) (35). Ub binds to DUBs with low affinity but high specificity, and thus mutations can be introduced to improve binding affinity without compromising the specificity. Indeed, UbVs for USP-family (34, 36) and OTU-family DUBs (34) can block the binding of substrate Ub to inhibit DUB activity.

We conducted five rounds of selections from an M13 bacteriophage pool representing the UbV library — Library 2 described previously (34) — for phage clones that bind to the biotinylated human full-length STAMBP protein (residues 1–424) (**Fig. 1A**). A total of 96 clones (48 from Round 4 and 48 from Round 5) were tested for binding activity to STAMBP using phage enzyme-linked immunosorbent assay (ELISA). Among them, 53 clones that displayed specific binding with STAMBP were subjected to DNA sequencing, which returned 14 unique UbV sequences (**Fig. S1**). We then selected the three most potent UbVs (denoted UbV^SP.1^, UbV^SP.2^, and UbV^SP.3^, where “SP” stands for STAMBP) for follow-up characterization (sequences shown in **Fig. 1B**). Finally, the binding specificity of these UbVs against a panel of 7 human DUBs of different families was assessed by phage ELISA (**Fig. 1C**). All three UbVs were confirmed to bind both the full-length and the JAMM domain of STAMBP (STAMBP^JAMM^), suggesting that the binding was mediated by the JAMM domain. In addition, UbV^SP.1^ exhibited cross-reactivity with STAMBPL1, while the other two variants showed significant preferential binding to STAMBP over STAMBPL1. Importantly, all three UbVs showed no binding to two USP-family DUBs (USP7 and USP14), a JAMM-family DUB complex BRISC, or an OTU-family DUB OTUD1.

**Figure 1.**
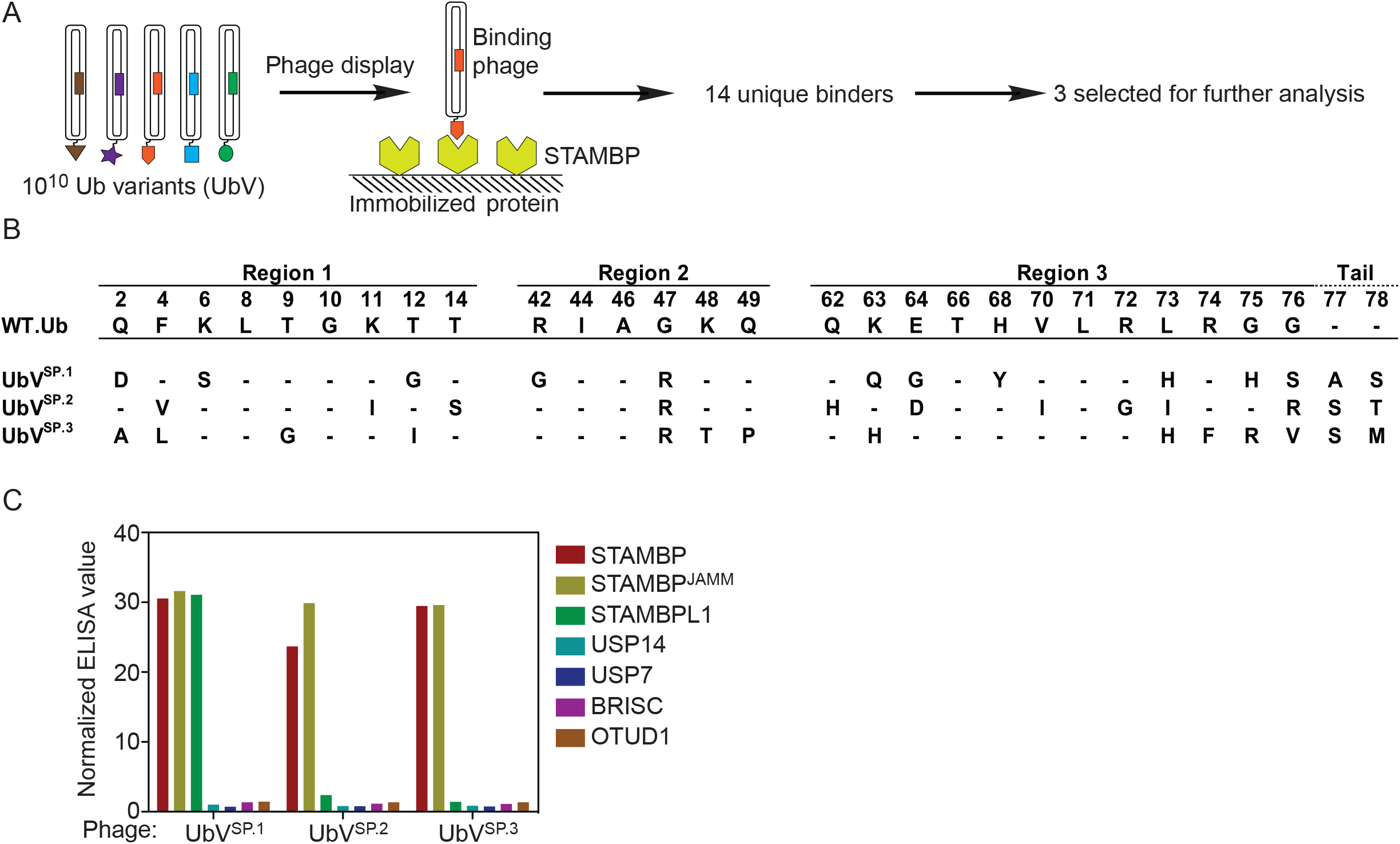
Ub variant (UbV) binders identified for STAMBP. **(A)** Schematic representation of the selection process. Within the UbV library, each phage particle displays a unique UbV. Binding phage were captured with immobilized STAMBP full length protein. After a total of five rounds of phage display selection, individual binding clones were subjected to sequencing. A total of 14 unique binders were identified, 3 of which were selected for downstream analysis. **(B)** Protein sequences of the three STAMBP UbVs (UbV^SP.1^, UbV^SP.2^, and UbV^SP.3^). Only the substitutions across the randomization surface of wild type Ub (WT.Ub) are shown. It should be noted that UbVs have two amino acid extensions at the C-terminal (position 77 and 78). Dashes indicate conservation of the WT.Ub sequence. **(C)** The binding specificities of phage-displayed UbVs were shown across a group of 7 deubiquitinases, as assessed by phage ELISA. Sub-saturating concentrations of UbV-phage were added to immobilized proteins as indicated. Bound phage were detected by the addition of anti-M13-HRP and colorimetric development of TMB peroxidase substrate. The mean value of absorbance at 450 nm was normalized to BSA control.

### Structural characterization of STAMBPL1 in complex with UbV^SP.1^

We then conducted crystallization trials for STAMBP^JAMM^ or STAMBPL1^JAMM^ in complex with the UbVs. Diffraction-quality crystals could only be obtained for STAMBPL1^JAMM^ in complex with the dual-specific UbV^SP.1^ despite a sequence identity of 68% between the two catalytic domains (**Fig. 2A**). We solved the crystal structure of the UbV^SP.1^/STAMBPL1^JAMM^ complex at 2.0 Å resolution (PDB:7L97, **Fig. 2B**, **Table 1**) in space group *C*2. The previous STAMBPL1^JAMM^/K63-diUb cocrystal structure (32) revealed the distal Ub has more extensive contact with the STAMBPL1 catalytic domain than the proximal Ub. Not surprisingly, UbV^SP.1^ occupies the distal Ub position in the structure. A total of 11 mutated residues differentiate UbV^SP.1^ from Ub.wt (**Fig. 1B**). The last two C-terminal residues (Ala77 and Ser78) of UbV^SP.1^ were not observed in the electron density map and were not modelled. Based on the crystal structure, it is evident that seven of the mutated residues (sites 2, 6, 12, 47, 63, 64, and 68) are distant from STAMBPL1 and unlikely to contribute to the tight binding observed. (**Fig. 2B**) The primary interaction interface was observed in the C-terminal tail of UbV^SP.1^, which includes three mutations, L73H, G75H, and G76S (**Fig. 2B**). PISA analysis (37) of the interface of the complex structure reveals the UbV has a buried surface area (BSA) of 1239 Å*2*. In contrast, the seven tail residues of the UbV (residues 70–76) contribute almost half of the BSA with 556 Å^2^.

**Figure 2.**
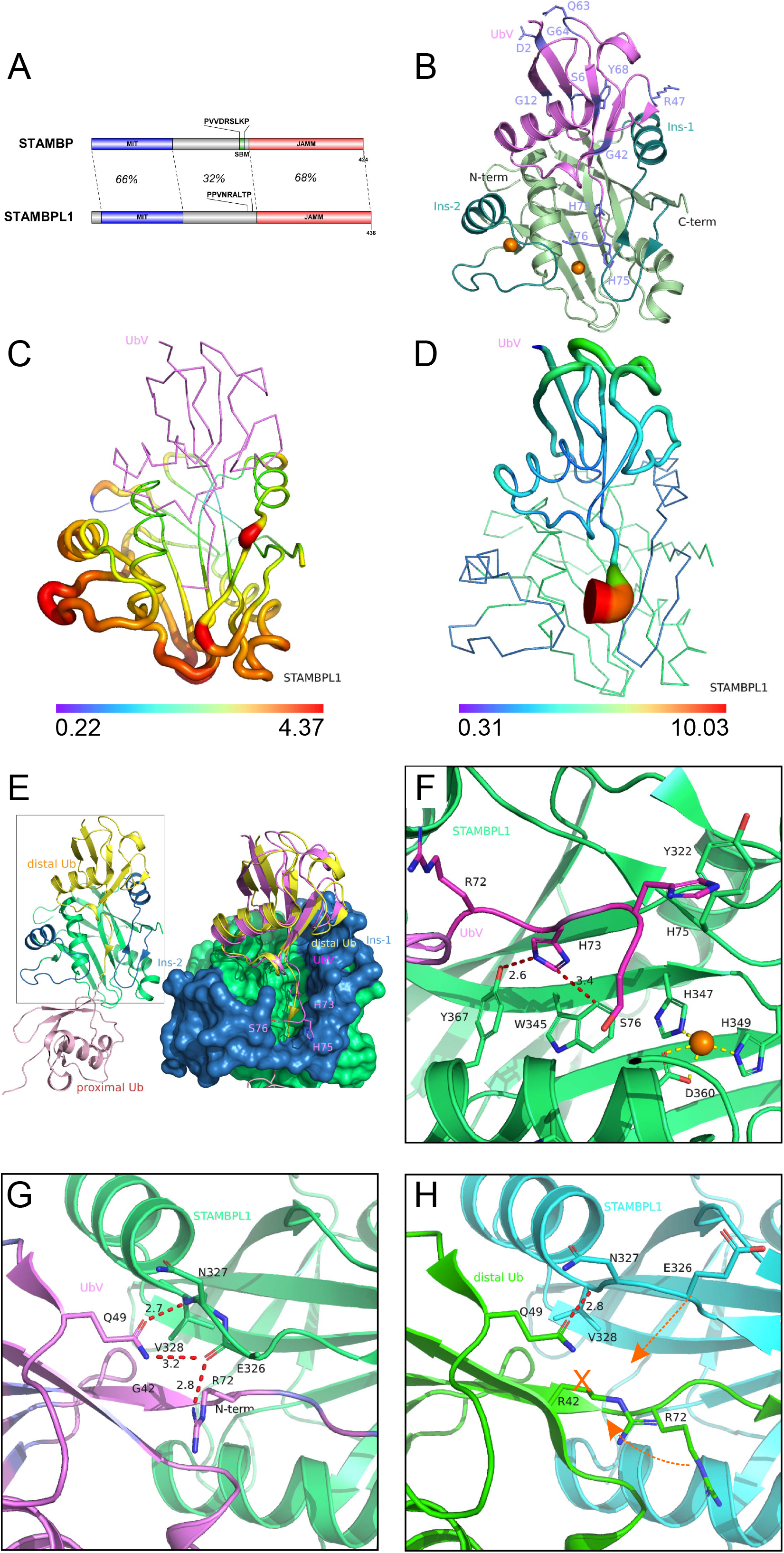
Crystal structure of the STAMBPL1-UbV^SP.1^ complex. **(A)** Domain architecture of STAMBP and STAMBPL1. The sequences corresponding to the SBM (SH3-binding motif) of STAMBP are indicated. **(B)** Overview of the complex structure. UbV^SP.1^ is shown in pink with variant residues shown as sticks and labelled. STAMBPL1 is shown in green, the two insertions (ins-1 and ins-2) in teal, and the two zinc ions are shown as orange spheres. **(C)** Conformational difference of the STAMBL1 catalytic domain in complex with UbV^SP.1^ vs. with K63-diUb (PDB:2ZNV, diUb omitted for clarity) in backbone sausage view. The radius of the sausage is normalized to the per residue CαRMSD. (in Å) between the minimum and maximum values. **(D)** Similar to panel (C), a sausage representation was applied to UbV^SP.1^ according to per residue C_α_ RMSD. compared to the distal Ub of the STAMBPL1/K63-diUb complex structure, when the STAMBPL1 structures are superimposed. **(E)** A comparison of the diUb/STAMBPL1 complex structure (left) and UbV/STAMBPL1 structure overlaid with diUb (right). The core of STAMBPL1 is shown in green, and the Ins-1 and Ins-2 insertions are in blue. It is evident that the UbV^SP.1^ (magenta) binds to the JAMM domain in the same orientation as that of the distal Ub (yellow), but unlike the distal Ub, the C-terminal tail of UbV^SP.1^ is excluded from the catalytic groove of STAMBPL1. The side chains of the variant residues H73, H75, and S76 unique to UbV^SP.1^ are show in sticks **(F)** A close-up view of the interactions around the C-terminal tail of UbV^SP.1^. H73 is buried in an aromatic pocket of STAMBPL1 comprised of Y367 and W345 and the side chain forms a strong hydrogen bond with the hydroxyl group of Y367, and a π-π stacking with W345 of the STAMBPL1. H75 of UbV^SP.1^ forms a T-shaped π-π stacking through the C_δ_-H with Y322 of STAMBPL1. The catalytic zinc ion and their coordinating residues are shown in spheres and sticks, respectively. **(G)** A close-up view of the interactions around E326 in STAMBPL1 ins-1. The side chain of Q49 and the guanidinium group of R72 forms a hydrogen bond network with the main chain of E326 and V328 from STAMBPL1. Note: the atoms in the side chain of E326 beyond C_β_ is not visible in the electron density map. **(H)** For comparison, in the STAMBPL1/K63-diUb structure (PDB:2ZNV), the side chain of R42 (G42 in UbV^SP.1^) precludes the formation of the hydrogen bond network seen in panel (G), the E326 loop of STAMBPL1, and the R72 of Ub.wt are expelled from forming a tight interaction.

**Table 1.** Crystallography data and refinement statistics.

|  |  |
| --- | --- |
| *Data collection* |  |
| Wavelength (Å) | 0.97949 |
| Resolution | 50.00–2.00 (2.03–2.00)* |
| Space group | $C_2$ |
| Unit Cell |  |
| $a, b, c$ (Å) | 63.71, 68.88, 57.33 |
| $\alpha, \beta, \gamma$ (°) | 90.0, 98.5, 90.0 |
| Measured reflections | 70314 |
| Unique reflections | 16078 |
| $I/\sigma I$ | 16.5 (1.8) |
| $R_{\text{merge}}$(%) | 12.3 (70.0) |
| $CC_{1/2}$ | 0.934 (0.769) |
| Completeness (%) | 99 (96.4) |
| Redundancy | 4.40 (3.40) |
| *Refinement* |  |
| Resolution (Å) | 50.00–2.00 |
| No. Reflections (test set) | 15388 (690) |
| $R_{\text{work}}/R_{\text{free}}$ (%) | 18.5/21.4 |
| No. Atoms |  |
| Protein | 1957 |
| Zinc | 2 |
| Water | 61 |
| B-factors (Å<sup>2</sup>) |  |
| Protein | 37.8 |
| Zinc | 40.3 |
| Water | 40.5 |
| RMSD |  |
| Bond lengths (Å) | 0.010 |
| Bond angles (°) | 1.377 |
| Ramachandran plot % residues |  |
| Favored | 98.8 |
| Allowed | 1.2 |
| Disallowed | 0.0 |

\*: values in parentheses are for highest resolution shell unless otherwise indicated.

The overall structure and mode of UbV^SP.1^ binding are similar to that of the distal Ub observed in the K63-diUb cocrystal structure. The RMSD of the backbone C_α_ atoms of STAMBPL1 in the two structures is only 0.29 Å over 146 residues, whereas the RMSD of UbV^SP.1^ compared to the distal Ub is 0.34 Å over 66 residues. Superposition of the complex structures in their entirety presents an overall RMSD of 0.56 Å over 212 residues, larger than the RMSDs of the individual molecules, suggesting a relative conformational movement. The conformational changes are most evident for three regions: a loop N-terminal to ins-1, the loops around the catalytic site, and the loops chelating the structural zinc ion. A CαRMSD-based sausage representation of the STAMBPL1 structure visualizes the regions that have the largest conformational change upon UbV^SP.1^ binding (**Fig. 2C**).

A comparison of the conformation of the UbV^SP.1^ and the distal Ub in the STAMBPL1/K63-diUb complex structure when the STAMBPL1^JAMM^ domain of the two structures are aligned reveals a significant conformational difference of the tails of the two bound Ub molecules (residues Arg72 and onwards; **Fig. 2D–G**). The backbone of the C-terminal tail of UbV^SP.1^ has the largest conformational movement compared to Ub.wt (**Fig. 2D**) and is excluded from the catalytic groove and projects in the direction of ins-2 while interacting with ins-1 (**Fig. 2E**). In contrast, the isopeptide bond of the physiological K63-diUb substrate is, as expected, buried in the catalytic groove and spans the catalytic site in an orientation that is primed for hydrolysis (**Fig. 2E, 2H**). Sequence comparison of UbV^SP.1^ with Ub.wt (**Fig. 1B**) and structural analysis (**Fig. 2F**) suggests the G75H mutation creates a steric hindrance that prevents the access of the tail to the catalytic groove, thereby providing a structural basis for the observed conformation. Instead, His75 forms a T-shaped π-stacking interaction with Tyr322 on ins-1 to stabilize the interaction. Meanwhile, the side chain of His73 in UbV^SP.1^ (equivalent to Leu73 in Ub.wt) forms a hydrogen bond with the hydroxyl group of Tyr367 and is buried in a hydrophobic pocket formed by Tyr367 and Trp345 of STAMBPL1. PISA analysis (37) also reveals that His73 has the largest BSA of 168 Å^2^ among all UbV^SP.1^ residues.(**Fig. 2F**).

Despite the ins-1 helix being a key region interacting with the distal Ub, it shows a relatively small conformational change compared to the rest of the molecule except for Glu326, a loop residue N-terminal to ins-1. Glu326 has the largest movement of C_α_ among the whole STAMBPL1^JAMM^ molecule at 4.4 Å (**Fig. 2C**). This prompted us to look further into the detailed interactions in this region. UbV^SP.1^ has a glycine at residue 42 (**Fig. 2G**), and the equivalent site on Ub.wt is occupied by an arginine (Arg42), with its side chain extending towards ins-1, effectively expelling the side chain of Glu326 away from the Ub.wt (**Fig. 2H**). In UbV^SP.1^, the R42G mutation creates space to accommodate the side chain of Arg72 in the C-terminal tail from UbV^SP.1^ (**Fig. 2G**) to flip towards this vacancy (**Fig. 2H**). A strong hydrogen bond network is then formed between the Arg72 guanidinium and the backbone carbonyl group of Glu326 in ins-1, and between the amide group of Gln49 of UbV^SP.1^ with the backbone carbonyl and amide of Val328 and Glu326, thus stabilizing the interaction between the STAMBPL1 and UbV^SP.1^ (**Fig. 2G**). In the STAMBPL1/K63-diUb complex, the side chain of Arg72 is oriented towards ins-2 but does not directly contact any STAMBPL1 residues (**Fig. 2H**). The coordinated mutation of R42G and flip of Arg72 side chain on UbV^SP.1^ with the movement of Glu326 on STAMBPL1 (**Fig. 2G–2H**) creates another hotspot for the tight interaction between UbV^SP.1^ and STAMBPL1.

### Mutational analysis identified hotspots of the tight binding between UbVs and STAMBP

To quantify the interaction of the UbVs with STAMBP and STAMBPL1, and to validate the key residues responsible for the tight binding, we used isothermal titration calorimetry (ITC) to measure the interaction thermodynamics (**Fig. 3**). All three UbVs evaluated showed a sub-µM affinity with STAMBP^JAMM^, and only UbV^SP.1^ showed a sub-µM affinity for STAMBPL1; Ub.wt does not show observable binding with STAMBPL1. While other UbVs had at least 14-fold higher affinity for STAMBP over STAMBPL1, UbV^SP.1^ has only 3.6-fold greater affinity towards STAMBP over STAMBPL1, indicating that it is a dual-specific binder. This is consistent with the result from phage ELISA (**Fig. 1C**).

**Figure 3.**
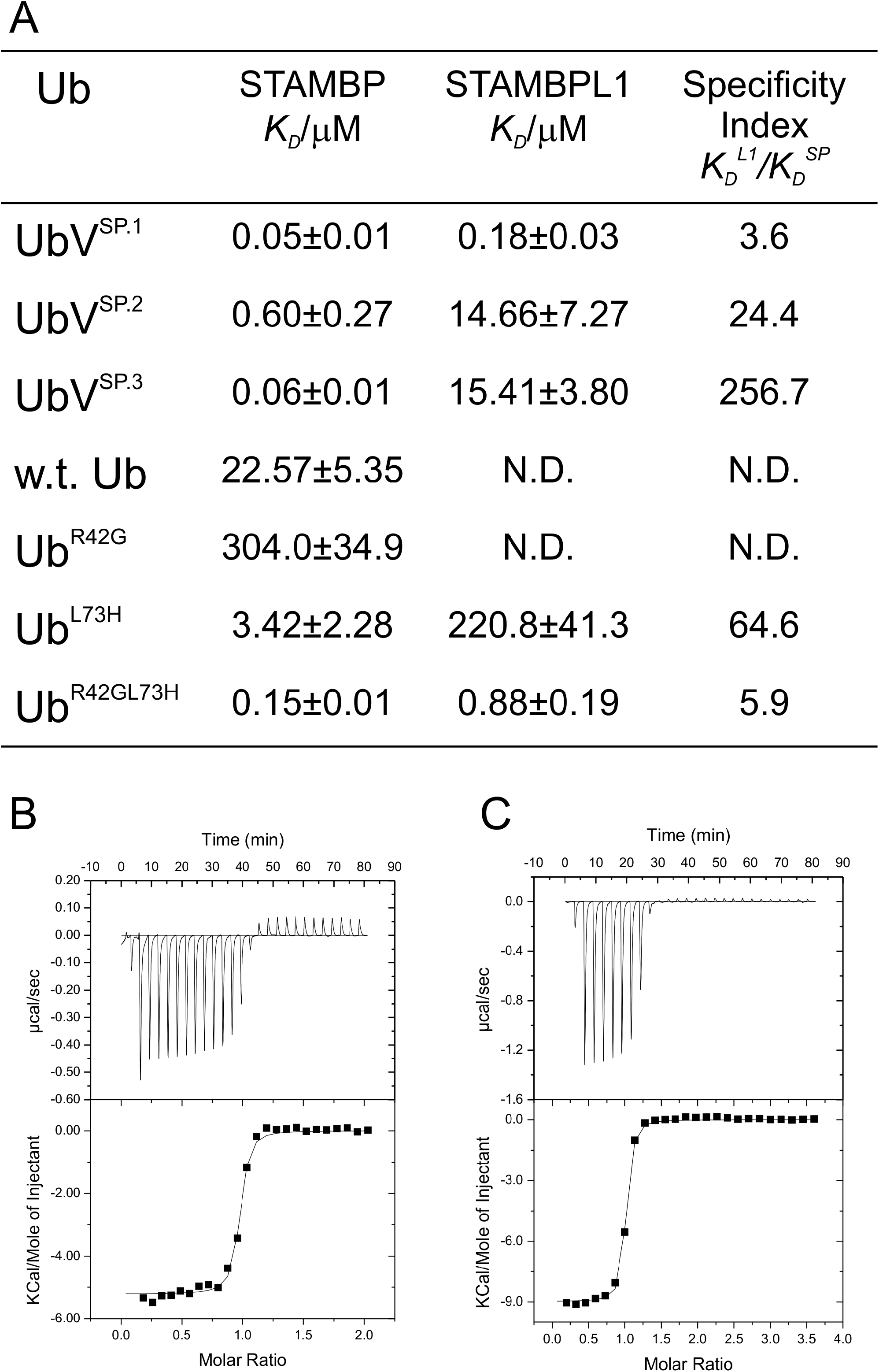
Isothermal titration calorimetry (ITC) measurement of the interaction between Ub, UbVs and STAMBP, STAMBL1. **(A)** Summary table of ITC measurements. N.D. stands for not determined, i.e. cannot be measured due to lack of interaction. **(B)** Thermodiagram of ITC measurement of STAMBPL1 with UbV^SP.1^. **(C)**Thermodiagram of ITC measurement of STAMBP with UbV^SP.1^. Specificity index is defined as the quotient of the dissociation constant of a UbV-STAMBPL1 complex divided by that of the same UbV in complex with STAMBP. The larger the value, the more specific the UbV is towards STAMBP.

Based on structural analysis, we hypothesized that R42G and L73H, among the 11 mutations of UbV^SP.1^, are the key residues that enhance the binding affinity of the UbV^SP.1^. To test this hypothesis, we constructed three mutants of Ub.wt, with either a single mutation Ub^R42G^, Ub^L73H^, or a double mutation Ub^R42GL73H^. We compared the binding affinity of these mutants with both STAMBP^JAMM^ and STAMBPL1^JAMM^ using ITC (**Fig. 3A–C**). Both Ub.wt and Ub^R42G^ showed no binding to STAMBPL1^JAMM^, while Ub^L73H^ showed only weak binding to STAMBPL1^JAMM^. The double mutant Ub^R42GL73H^, however, showed a sub-μM affinity (*K*_*D*_ of 0.88 μM) only 5-fold weaker than that of UbV^SP.1^, suggesting a synergistic effect of the two mutations. The binding of the mutants with STAMBP^JAMM^ show a similar trend; the R42G mutation reduced the binding of the Ub to STAMBP^JAMM^ and the L73H mutation only slightly increased the affinity. The double mutant Ub^R42GL73H^, however, resulted in a sub-μM affinity at 0.15 μM, only 3-fold weaker than the UbV^SP.1^. These results suggest the mutations at sites 42 and 73 are the hotspots for UbV^SP.1^ for its high binding affinity towards both STAMBP^JAMM^ and STAMBPL1^JAMM^.

### UbV^SP.1^ and UbV^SP.3^ are potent STAMBP inhibitors

To investigate the inhibitory effects of UbVs on STAMBP *in vitro*, we first performed a fluorescence resonance energy transfer (FRET)-based K63-diUb substrate cleavage assay. We found that UbV^SP.1^ and UbV^SP.3^ are potent inhibitors for the isopeptidase activity of STAMBP^JAMM^. The half-maximal inhibition (IC_50_) was 8.4 nM for UbV^SP.1^ and 9.8 nM for UbV^SP.3^ both having a Hill slope of −0.8 (**Fig. 4A–B**). To benchmark the inhibitory potency of our UbV inhibitors, we obtained previously reported STAMBP small-molecule inhibitors having two different chemistries each reported as BC-1471 (33): consisting of a core 2-6-morpholino-4-oxo-3-phenethyl-3,4-dihydroquinazolin-2-yl-thio- and an *N*-linked tetrahydrofuran-2-yl-methyl acetamide (CAS 896683-84-4, racemate) or a furan-2-yl-methyl-acetamide (CAS 896683-78-6). In both cases, we did not observe the inhibitory activity reported for BC-1471 as there was no difference in the diUb deconjugation reaction rate of STAMBP^JAMM^ with or without the compounds (**Table 2**).

**Figure 4.**
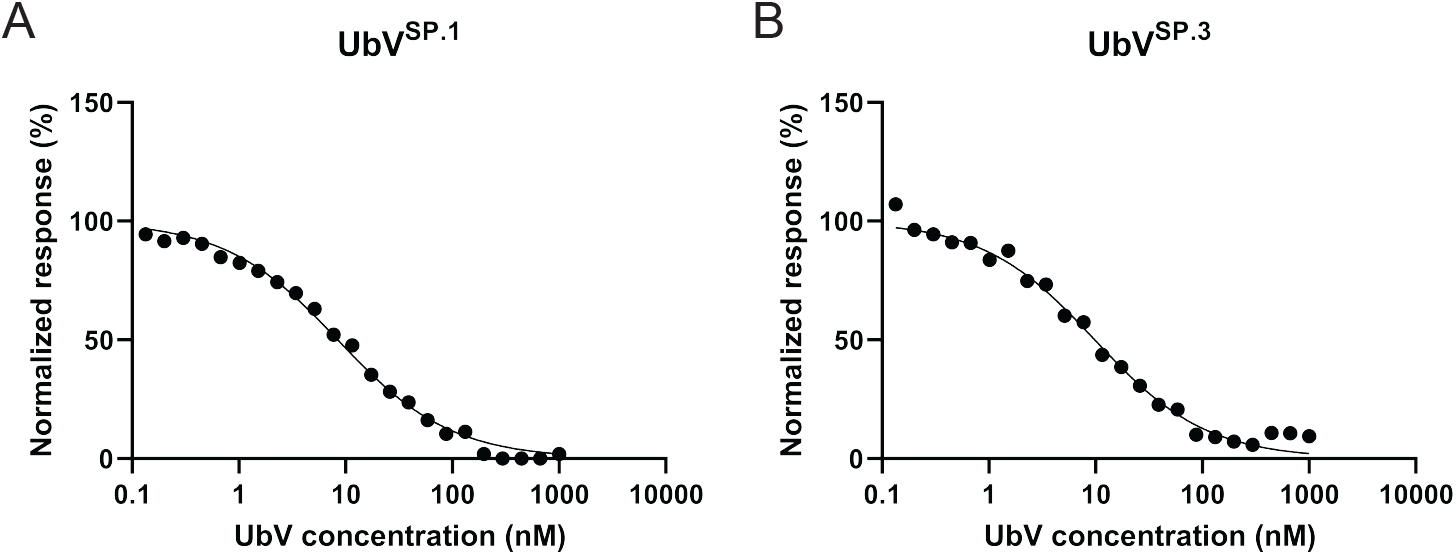
UbV inhibition of STAMBP^JAMM^ diUb isopeptidase activity. Inhibition of STAMBP^JAMM^ isopeptidase activity by **(A)** UbV^SP.1^ and **(B)** UbV^SP.3^ shown as dose-response curves using a K63-linked diUb FRET substrate. The IC_50_ value was determined as the concentration of UbV that inhibited 50% of proteolytic activity. Curves were fit by non-linear regression using the formula for inhibitor concentration vs. normalized response (variable slope) in GraphPad Prism 8. A Hill slope of −0.8 was calculated for both inhibitors representing a single-site binding model.

**Table 2.** FRET based diUb cleavage assay to measure deubiquitinating activity.

| Enzyme | Activity with 1 μM inhibitor<br>(% of uninhibited activity)* |
| --- | --- |
| STAMBP/STAM1+UbV<sup>SP.1</sup> | 7.2±0.6 |
| STAMBP/STAM1+UbV<sup>SP.3</sup> | 13.4±0.2 |
| STAMBPL1+UbV<sup>SP.1</sup> | 44±3 |
| STAMBPL1+UbV<sup>SP.3</sup> | 118±5 |
| STAMBP<sup>JAMM</sup>+UbV<sup>SP.1</sup> | 0 |
| STAMBP<sup>JAMM</sup>+UbV<sup>SP.3</sup> | 11±2 |
| STAMBP<sup>JAMM</sup>+BC-1471<br>(CAS 896683-84-4) | 107±2 |
| STAMBP<sup>JAMM</sup>+BC-1471<br>(CAS 896683-78-6) | 96±3 |

\* Reactions were measured in triplicate at room temperature with 5 nM enzyme and 100 nM diUb K63 TAMRA substrate. Reported activity is normalized to identical reactions without added inhibitor.

Next, we compared the K63-diUb (FRET) cleavage inhibition by the UbVs with STAMBP^JAMM^ to the inhibition of full-length STAMBP activated by STAM protein and full-length STAMBPL1. STAM contains three Ub-binding domains: VPS27/Hrs/STAM (VHS), Ub-interacting motif (UIM), and SH3 domains. The SH3 domain mediates the interaction of STAM with the SBM motif N-terminal to the JAMM domain of STAMBP. The VHS domain shifts the cleavage preference of STAMBP to longer poly-Ub chains by binding to a Ub that is spatially distant from the JAMM domain and helps position the cleavage site (38), while the UIM domain is proposed to bind to the proximal Ub when the JAMM domain of STAMBP recognizes the distal Ub of diUb (39, 40). As expected from IC_50_ assays, UbV^SP.1^ resulted in complete inhibition of STAMBP^JAMM^ and UbV^SP.3^ reduced the activity to 10% of the uninhibited enzyme (**Table 2**, UbV concentration=1 μM). Full-length STAMBP activated with STAM1 was inhibited by both UbV^SP.1^ and UbV^SP.3^, and the activity was reduced to 7% and 13%, respectively. The activity of the paralog STAMBPL1 with the K63-diUb FRET substrate was inhibited to 44% by UbV^SP.1^; however, there was no observable inhibition by UbV^SP.3^. This is consistent with the binding affinity data shown in **Fig. 3**.

Next, we assessed the effect of UbV^SP.1^ and UbV^SP.3^ using a K63-linkage poly-Ub (Ub_2_-Ub_7_) cleavage assay. In this assay, the isopeptidase activity of STAMBP can be visualized by monitoring the appearance of mono-Ub (Ub_1_) and the disappearance of the Ub_2_-Ub_7_ bands (**Fig. 5A**). STAMBP in complex with different UbV inhibitors were incubated with K63-linkage poly-Ub, and STAMBP^JAMM^ samples in the presence and absence of Ub.wt were used as controls. Both UbV^SP.1^ and UbV^SP.3^ potently inhibited the isopeptidase activity of STAMBP^JAMM^ (**Fig. 5B**) and full-length STAMBP (**Fig. 5C**) towards the cleavage of K63-linked poly-Ub chains. In addition, UbV^SP.1^ appeared to inhibit STAMBPL1 to some extent, whereas UbV^SP.3^ showed no inhibitory effect with STAMBPL1 (**Fig. S2A**).

**Figure 5.**
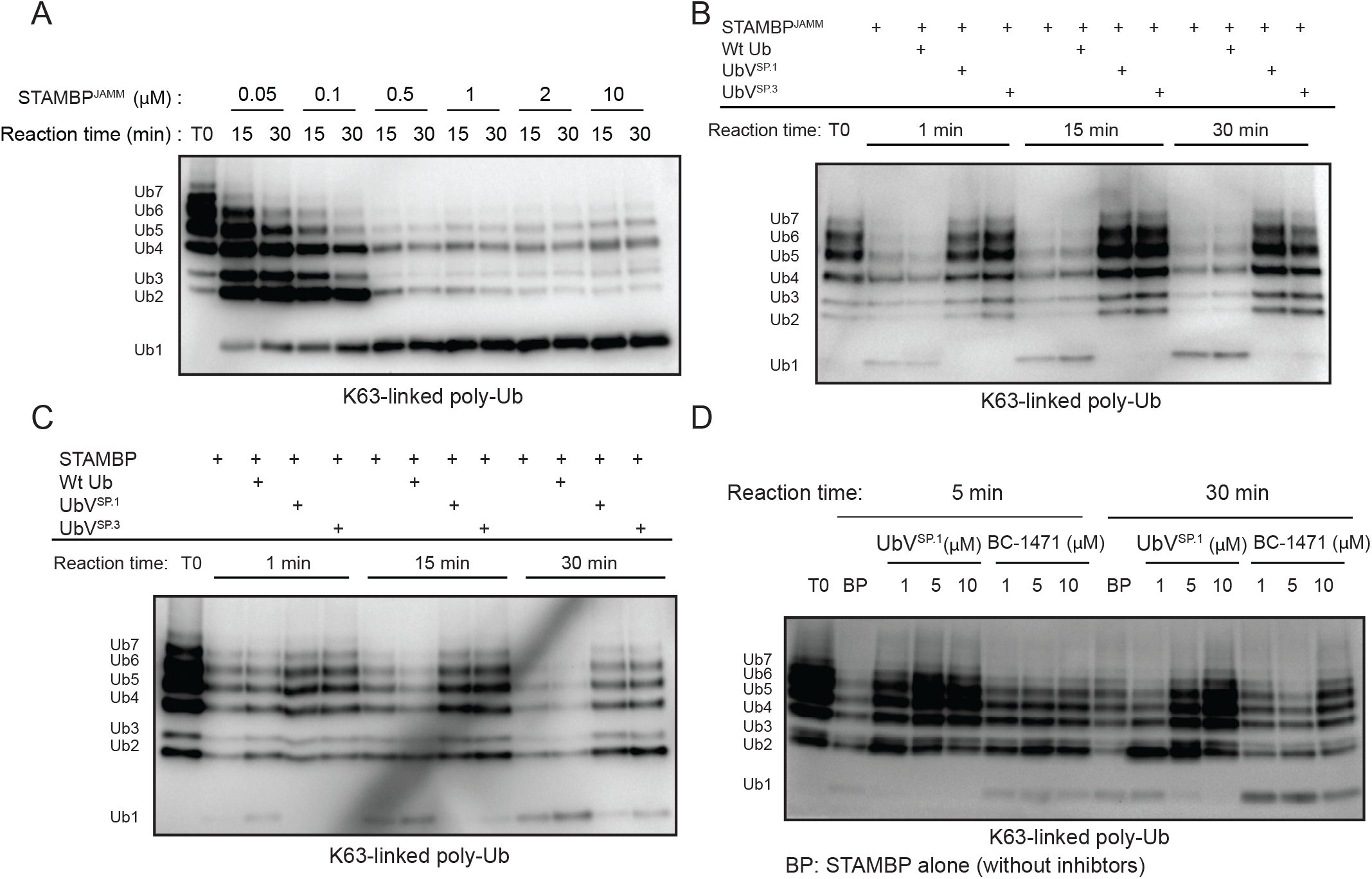
UbVs are potent inhibitors of STAMBP. **(A)** Assessment of deubiquitination activity of STAMBPJAMM using a biotinylated K63-linked poly-Ub (Ub_2_-Ub_7_) substrate. Purified STAMBP^JAMM^ proteins with different concentrations as indicated were incubated with the poly-Ub substrate at 37°C for a time course of 30 minutes. Samples were resolved by SDS-PAGE and visualized by western blotting, which was probed with ExtrAvidin-HRP (EA-HRP) to detect biotin-Ub. Deconjugation of poly-Ub chains was observed by the disappearance of Ub2-Ub7 and appearance of the digestion product mono-Ub (Ub_1_). **(B-C)** Inhibition of STAMBP^JAMM^ **(B)** and full-length STAMBP **(C)** deubiquitination activity by UbVs. As in **(A)**, purified DUB proteins were incubated with the indicated UbV or Ub.wt (negative control) and poly-Ub substrate at 37°C for a time course of 30 minutes. Inhibition of isopeptidase activity was indicated by the retention of Ub_2_-Ub_7_ and reduced appearance of the digestion product mono-Ub (Ub_1_). **(D)** Comparison of inhibitory efficacy of UbV^SP.1^ to previously published STAMBP small molecule inhibitor BC-1471 against isopeptidase activity of full-length STAMBP (BP). Experiments were conducted as described above.

We then compared the inhibitory effects to the small-molecule inhibitor BC-1471 (33), UbV^SP.1^ exhibited a much greater inhibitory effect to STAMBP than BC-1471 (**Fig. 5D**). Importantly, the inhibition by UbV^SP.1^ is dose-dependent, but this is not the case for BC-1471 (**Fig. S2B**), further confirming that BC-1471 is unlikely a potent inhibitor of STAMBP. In addition, we observed inhibition of STAM1-activated full-length STAMBP by both UbV^SP.1^ and UbV^SP.3^ in this chain cleavage assay (**Fig. S2C**). Finally, neither UbV^SP.1^ and UbV^SP.3^ showed inhibitory effects to other JAMM DUBs, such as BRISC (**Fig. S2D**), and MYSM1 (**Fig. S2E**), which indicated the inhibitory specificity of UbV^SP.1^ and UbV^SP.3^.

## Discussion

Over the past few years, the phage-display based protein engineering platform targeting Ub-mediated protein-protein interactions (41) has been utilized to develop potent and specific modulators for a repertoire of UPS components, including E2 conjugating enzymes (42), E3 ligases (43–45), and DUBs (36, 46–48). These DUBs include seven USPs (USP2, USP7, USP8, USP10, USP15, USP21, and USP37); one OTU (OTUB1); and two viral papain-like proteases (MERS-CoV PLpro and CCHFV-L, which have USP and OTU structural fold, respectively) (36, 41, 46–48).

The UbVs described herein for STAMBP and STAMBPL1 are the first examples of protein-based potent and specific deubiquitination inhibitor developed for the JAMM family DUBs. Previously, a moderately selective (>5-fold towards several JAMM DUBs) RPN11 inhibitor capzimin was generated by optimization of a non-specific small molecule inhibitor 8-TQ, which has structural similarity with chelating agent 8-hydroxyquinoline (25). Moreover, a small-molecule inhibitor CSN5i-3 was developed to inhibit CSN5, demonstrating the feasibility of targeted inhibition of JAMM DUBs (26). CSN5 is unique in that it needs the whole CSN complex to elicit deneddylation activity and has NEDD8-conjugated cullin-RING E3 ligases as specific substrates. While BC-1471 was recently reported as a STAMBP chemical inhibitor (33), we were not able to detect its inhibitory activity in our *in vitro* assay. We note that BC-1471 was identified from an *in silico* screening method with no confirmation using a cocrystal structure or an orthogonal biophysical technique to validate a direct binding (33). In addition, previous *in vitro* assays did not show complete inhibition of STAMBP activity by BC-1471 at high concentrations (100 μM) or under a native cellular phenotype such that the observed inhibition could be the result of off-target effects (33).

Similar to other UbVs targeting USPs, including UbV^core^ for USP37, UbV^7.2^ for USP7, and UbV^10.1^ for USP10 (36, 46), UbV^SP.1^ has enhanced affinity primarily due to amino-acid substitutions at the C-terminal variable region (N60−G76+). Structural characterization of the STAMBPL1^JAMM^-UbV^SP.1^ complex, along with mutation analysis of amino-acid substitutions, identified hotspot residues contributing to the enhanced affinity of the variants for the catalytic domains of STAMBP and STAMBPL1. With a single residue substitution (L73H) in the C-terminus alone, the affinity of Ub.wt for STAMBP was enhanced nearly 7-fold. Subsequently, a double mutation (R42G, L73H) enhanced the affinity of Ub.wt for STAMBP by 150-fold.

The simplicity of the phage display selection process opens the opportunity for rapidly developing high-affinity and selective DUB inhibitors. Another advantage of the UbV platform over the peptide or small molecule compound screening strategies comes from the fact that Ub.wt is a low-affinity natural substrate for the target DUBs, and the variable C-terminal region are topographically placed in the proximity of the catalytic groove (49). Theoretically, this confers a significant reduction of the search space to engage binding during the selection process, further improving the efficiency in selecting high-affinity binders. In addition, the large Ub binding surface on the DUBs provides opportunities for the UbV to bind to the cognate DUB and compete with substrate Ub.wt without direct interaction with the catalytic site residues. For example, the C-terminal tail of UbV^8.2^ for USP8 (34) binds to a cleft between the blocking loop and the fingers subdomain that is remote from the catalytic groove yet still compete with Ub.wt binding. For reference, UbV^8.2^ is rotated about 40° when compared to the binding mode of Ub.wt in other USP complexes (47). Therefore, targeting the substrate Ub.wt binding region of the JAMM domain by UbVs is novel compared to current non-specific chemical reagents targeting metalloproteases through the chelation of the catalytic zinc ion.

Furthermore, improved affinity and specificity has been achieved for several UbVs (e.g., UbV^15.1a–e^ for USP15 and UbV^Fl11.1^ for SKP1-FBL11 complex) through additional residue insertions in the β1-β2 loop, extending the interaction surface of the UbV (45, 47, 50). The UbVs developed for STAMBP and STAMBPL1 have multiple mutations in β1, β2, and the loop they encompass; however, these mutations have little effect on UbV binding due to the orientation of distal Ub binding to the JAMM domain. The β1-β2 loop is poised to make substantial interactions with β-strands 2, 3, and 6 of STAMBPL1 with a similar peptide insertion as was employed previously for USP15 and SKP1-FBL11. Thus, specificity and affinity could potentially be improved with insertions in the β1-β2 loop and can be explored in the future.

Intriguingly, UbV^SP.1^ has nanomolar affinities for both STAMBP and the paralog STAMBPL1, while UbV^SP.3^ has a similar affinity for STAMBP but a reduced affinity for STAMBPL1. Disparity in the affinities of the two UbVs was also translated to the inhibitory effects of the respective UbVs on isopeptidase activity. UbV^SP.1^ was equally effective at inhibiting STAMBP and STAMBL1 activity, whereas UbV^SP.3^ had no apparent effect on STAMBPL1 activity. Structural and sequence similarities of STAMBP and STAMBPL1 suggest that UbV^SP.1^ likely binds in a similar manner to both enzymes. The noticeably stronger affinity (3-fold, **Fig. 3A**) of UbV^SP.1^ for STAMBP is likely due to the substitution of the Val328 (STAMBPL1) in the ins-1 loop by a glutamic acid (Glu316 in STAMBP), which may introduce an additional salt bridge with Arg72 from UbV^SP.1^ (**Fig. S3A**). Of the two hotspot mutations (R42G and L73H) identified in UbV^SP.1^, UbV^SP.3^ also harbors the L73H mutation but preserves the Arg42 of Ub.wt (**Fig. 1B**). The weaker affinity of UbV^SP.3^ for STAMBPL1 and the resulting lack of inhibition of STAMBPL1 activity may be explained by the difference in the interaction with these residues. The preservation of Arg42 in UbV^SP.3^ prevents its Arg72 from flipping towards ins-1 and participating in the hydrogen bond network as observed in the STAMBPL1:UbV^SP.1^ structure. In contrast, the side chain of Glu316 of STAMBP could form water-bridged hydrogen bonds with Arg42 and Arg72, thereby resulting in a reorganization of the ins-1 loop. Although no structure of human STAMBP in complex with Ub.wt has been reported, such a water-bridged interaction was observed in the complex structure of the yeast STAMBP orthologue Sst2 in complex with K63-diUb (PDB:4NQL, **Fig. S3B**). The increased affinity of UbV^SP.3^ for STAMBP over Ub.wt is partially explained by the same L73H substitution as in UbV^SP.1^ (as demonstrated from the observed 7-fold increase in affinity from ITC experiments). Additional affinity could be caused by additional hydrophobic interactions of the introduced R74F mutation with residues Tyr310, Phe395, and Val347 (Tyr322, Phe407, and Val359 STAMBPL1 numbering). The five residues of UbV^SP.3^ tail (Phe74–Met78) differ significantly from those in UbV^SP.1^ (**Fig. 1A**) and may form additional enzyme-specific interactions divergent from those observed for the respective residues in the STAMBPL1:UbV^SP.1^ structure. While a relative specificity is observed for UbV^SP.1^ and UbV^SP.3^ against other DUBs, a thorough proteomics analysis may be required before using these molecules in target validation experiments. For example, there are more than 20 different Ub-binding domains (UBDs) identified to date and they are quite diverse at the structure level (51). We noted that STAMBP UbVs do not have the key mutation at position 10 (i.e. G10A/V mutation that can lead to UbV dimerization) necessary to increase affinity for binding to Ub-interacting motifs (UIMs) (46, 52, 53). However, UbV^SP.1^ and UbV^SP.3^ may potentially interfere with certain Ub-UBD interactions. Like the off-target effects in drug discovery, whether UbV^SP.1^ or UbV^SP.3^ may bind to other proteins, including those do not contain UBDs, should be tested in separate cellular experiments.

Given the success of generating first-in-class UbV inhibitors for STAMBP and STAMBPL1, further development of UbV modulators is conceivable for the remaining 13 members of the JAMM family, including the 7 pseudo-DUBs. While the structural elements of the JAMM domain persist, many of the residues composing the canonical Ub binding site are divergent. Employing UbV selection strategies opens the opportunity of utilizing the deviation from the canonical binding site as a unique target for protein-protein interactions. Of particular interest are the BRCC36-ABRAXAS or -KIAA0157 complexes in the BRCA1-A and BRISC holocomplexes, respectively (54). The aforementioned heterodimers with BRCC36 form the catalytic cores of their respective holocomplexes, incorporating a dimer of JAMM and pseudo-JAMM domains. Targeting these structures with UbVs would provide a more rapid and specific option than conducting small molecule inhibitor screens. Finally, the recombinant UbV modulators of the pseudo-DUBs will be ideal probes for evaluating the biological functions of these pseudo-enzymes without ablating their potential roles as molecular scaffolds in cellular processes.

## Experimental procedures

### Protein overexpression and purification

All constructs for structural biology and isothermal titration calorimetry (ITC) binding measurement were cloned into a pET28-MHL vector, which encodes a TEV protease cleavable N-terminal His6-tag that will leave a non-native glycine residue after TEV treatment of the target protein, and overexpressed in *E. coli* strain BL21 (DE3) with 250 μM isopropyl-β-D-thiogalactopyronoside (IPTG) induction using a protocol described previously (55). 1×PBS with 5% glycerol was used as the base buffer for the whole purification process with added supplements in different purification steps. Briefly, cell pellets from 2 L culture were used for each construct. The sonication disrupted cells were first clarified with centrifugation and purified using open column Ni-NTA chromatography. The bound proteins were washed with the base buffer supplemented with 20 mM imidazole and eluted with the buffer containing 250 mM imidazole. The eluates were pooled and digested with TEV protease at 30:1 (w/w) protein:TEV, then dialyzed overnight against 1×PBS. The dialyzed solution was passed through 5 mL Ni-NTA resins again to remove uncut proteins and TEV protease. The flow-through was further purified by cation-exchange chromatography on a 5 mL HiTrap S column (GE Healthcare) with a salt gradient of 0 to 1 M NaCl in 1×PBS supplemented with 1 mM DTT. The fractions containing the target proteins were collected and loaded on a Superdex 200 size exclusion chromatography (SEC) column. The eluates were then collected, pooled, and concentrated by centrifugal filters (Amicon mwco. 10 kDa) to a final concentration of 10–20 mg/mL. The final purities of the proteins were about 99%, judging from SDS-PAGE and Coomassie Brilliant Blue (CBB) staining. The removal of the His6 tag of the proteins was confirmed using mass spectroscopy. Three constructs corresponding to the full-length (residues 1–424) and JAMM domain (residues 219–424) of STAMBP in reference to NP_998787.1, and the JAMM domain (residues 263–436) of STAMBPL1 in reference to NP_065850.1 were used for UbV selection. The constructs were cloned into a modified pET-28a vector encoding a TEV cleavable N-terminal avi-tag for *in vivo* biotinylation and a C-terminal His6 tag and transformed into a BL21 (DE3) *E. coli* strain harboring a plasmid containing the BirA ligase. The proteins were purified using Ni-NTA chromatography followed by SEC before being used for UbV selection. The purities of the preparations are higher than 95% based on SDS-PAGE and CBB staining.

### Crystallization of the protein complex

Different combinations of STAMBP^JAMM^ or STAMBPL1^JAMM^ in complex with UbV were prepared for crystallization trials. The corresponding pairs of two proteins were mixed at 1:1.1 molarity ratio (DUB:UbV) and loaded on a Superdex 200 SEC column. The fractions corresponding to the protein complex were pooled and concentrated by centrifugal filters (Amicon mwco. 30kDa) to a final concentration of 16 mg/mL. Crystallization trials were set using 0.5 *μ*L protein solution plus 0.5 *μ*L reservoir solution in a sitting drop vaporization setup. Three crystal screening kits were tried for each sample: SGC-II, Red Wings, and Hampton Research Protein Complex. Multiple crystal forms were obtained, but only certain conditions gave a crystal containing both polypeptides. For all the combinations of the DUB:UbV complexes tried, we were only able to obtain a crystal of the STAMBPL1^JAMM^/UbV^SP.1^ complex. The crystal used for data collection was optimized using hanging drop vaporization set up in a reservoir solution containing 20% PEG1500, 0.2 M NaCl, 0.1 M HEPES, pH 7.5. The crystal was transferred to a cryoprotectant containing 15% ethylene glycol in the reservoir solution before flash-freezing in liquid nitrogen.

### Diffraction data collection and structural determination

X-ray diffraction data for structure determination were collected at the Canadian Light Source Inc. (CLSI) beamline 08ID-1 (CMCF-ID) (56). The dataset was processed with the HKL-3000 suite (57). The structures were solved by molecular replacement using PHASER v.2.6.1 using PDBs 2ZNR and 1UBQ as the search template for STAMBPL1 and UbV^SP.1^, respectively. COOT was used for model building and visualization, REFMAC (v.5.8.0135) for restrained refinement. The final model was validated by MOLPROBITY. All molecular graphics were prepared with PyMOL v.2.4.0 (Schrödinger, Inc.)

### ITC measurement

For ITC measurement, proteins were dialyzed overnight with PBS buffer containing 1 mM TCEP and adjusted to a final concentration of 50 μM. All ITC measurements were performed at 25°C on a NanoITC (TA Instruments). A total of 25 injections, each of 5 μL Ub.wt, mutants, or UbVs were delivered into a sample cell of 200 μL containing one of the STAMBP^JAMM^ or STAMBPL1^JAMM^. The data were analyzed using the NanoAnalyzer software supplied by the manufacturer and fitted to a one-site binding model.

### Phage display selections and ELISA to evaluate binding

The phage-displayed UbV library used for selection was re-amplified from Library 2 as previously described (34). Protein immobilization and the following phage selections were done according to standard protocols (58). Briefly, purified STAMBP proteins (full-length or JAMM domain) were coated on 96-well MaxiSorp plates by adding 100 μL of 1 μM proteins and incubating overnight at 4°C. Five rounds of phage display selections were performed as follows: a) Preparation of the phage library pool, within which each phage particle displays a unique UbV and encapsulates the UbV encoding DNA in a phagemid; b) The pool of phage-displayed UbV library are applied to immobilized STAMBP; c) STAMBP-binding phage are captured, and non-binding phage are washed away; d) Bound phage are amplified by infection of bacterial host *E. coli* (OmniMAX); e) Individual phage with improved binding properties obtained from round 4 and round 5 are identified by phage ELISA (see below) and subjected to DNA sequencing of the phagemids to obtain UbV sequences (44). High-affinity UbVs identified from phage ELISA were cloned into expression vectors, including a FLAG-tag epitope, expressed in *E. coli*, and purified. For the ELISA experiments, proteins were immobilized on 384-well MaxiSorp plates (Thermo Scientific 12665347) by adding 30 μL of 1 μM protein solution for overnight incubation at 4ºC. Phage and protein ELISAs with immobilized proteins were performed as described before. Binding of phage or FLAG-tagged UbVs was detected using anti-M13-HRP (GE Healthcare 27942101) or anti-FLAG-HRP antibody (Sigma-Aldrich A8592), respectively. To measure the half-maximal effective binding concentration (EC_50_) of the UbVs’ binding to the immobilized DUB, the concentration of UbVs in solution was varied from 0 to 4 μM by two-fold serial dilutions. EC_50_ values were calculated using the GraphPad Prism (version 5.0a) software with the built-in equation (non-linear regression curve).

### Inhibition of STAMBP^JAMM^ diUb isopeptidase activity by UbVs

Inhibition concentrations were determined using a FRET-based diUb cleavage reaction to measure isopeptide hydrolysis by the STAMBP^JAMM^ domain. Fluorescence of hydrolyzed K63-linked diUb (TAMRA/QXL position 3 labeled, Boston Biochem #UF-330) was measured on a BioTek synergy LX plate reader equipped with a red filter cube assembly (ex. 530 nm, em. 590 nm) for 60 minutes at ambient temperature (22°C). Enzyme (2 nM) and UbVs were pre-incubated for 10 minutes at ambient temperature before addition of the diUb substrate (100 nM) in 50 μL reaction buffer (50 mM HEPES pH7.5, 100 mM NaCl, 0.01% v/v Tween 20, 0.01 mg/mL BSA, 5 mM DTT). Reactions were replicated in triplicate (N=3) for varying concentrations of UbV (1 μM to 0.1 nM). The rate of reaction was determined from the slope of the initial linear phase of the reaction. Reaction rates were averaged and normalized to the negative control (no inhibitor) then plotted against UbV concentrations in GraphPad Prism 8. 50% inhibition concentrations were determined by non-linear regression using the formula for inhibitor concentration vs. normalized response (variable slope).

### Comparison of diUb Isopeptidase Inhibition with Full-Length STAMBP and STAMBPL1

To compare the inhibition of STAMBP^JAMM^ by UbV^SP.1^ and UbV^SP.3^ with wild-type enzymes, inhibition was evaluated with full-length STAMBP (Boston Biochem #E-549) (activated with STAM1 (Boston Biochem #E-550)) and STAMBPL1 (Boston Biochem #E-551). Reactions were measured with similar conditions as were used for measuring STAMBP^JAMM^ inhibition. Each reaction instead had 1 μM of UbV and 5 nM of enzyme with 100 nM diUb (K63). STAMBP was pre-incubated with STAM1 (final concentration of 100 nM) for 10 minutes prior to the addition of UbV (final concentration of 1 μM). The BC-1471 compounds (TargetMol, CAS 896683-84-4 and CAS 896683-78-6) were also evaluated with STAMBP^JAMM^ under identical conditions with a final concentration of 1 μM. Initial reaction rates were measured from the slope of the linear portion of the reaction. Reactions were measured in triplicate (N=3) and averaged to determine the inhibited reaction rates. Inhibited reaction rates were normalized to control reactions for each enzyme without inhibitor.

### Inhibition assay with poly-Ub cleavage

Inhibition assays were conducted with biotinylated K63-linked poly-Ub chains, a mixture of poly-Ub chains (Ub_2_–Ub_7_) (Boston Biochem #UCB-330-050), as the substrate. Assays were performed in a buffer [50 mM HEPES (pH 8.0), 150 mM NaCl, 1 mM TCEP] containing 0.2 μM substrate, DUB (in sub-μM range with some variations), and Ub.wt. (in same concentration as UbVs), UbVs (UbV:DUB = 5:1 for STAMBP and STAMBP^JAMM^, 20:1 for STAMBPL1, 10:1 for BRISC and MYSM1), STAM1 (STAM1:DUB = 5:1) or BC-1471 (TargetMol) at indicated concentrations. Ub.wt, UbVs, and BC-1471 were incubated with DUB at room temperature for 5 min before adding the poly-Ub substrates, and STAM1 was added and pre-incubated with DUB before adding UbVs/ Ub.wt. Reaction mixtures were incubated at 37 °C for indicated reaction times and stopped by adding 1 mM EDTA and SDS-PAGE loading buffer. Samples were resolved by tricine-SDS-PAGE and visualized by Western-blots probed by Ub-antibody Extravidin^®^-Peroxidase (Sigma #E2886). All experiments were conducted in triplicate (N=3).

## Data availability

The atomic coordinates and structure factors (Code:7L97) have been deposited in the Protein Data Bank (https://wwpdb.org). PyMOL script for generating the sausage view is uploaded to GitHub at https://github.com/tongalumina/rmsdca. All other data are included in this manuscript.

## Acknowledgments

We greatly appreciate help from Dr. Sachdev S. Sidhu (University of Toronto) who provided the UbV library. We sincerely thank Dr. Elton Zeqiraj (University of Leeds) for providing BRISC complex proteins. Q.L. is supported by an OGS scholarship. E.M. was a MITACS Industrial Postdoctoral Fellow with funding provided by MITACS and ProteinQure Inc. WZ is currently a CIFAR Azrieli Global Scholar in the Humans & The Microbiome Program. W.Z. is also a recipient of the Cancer Research Society/ BMO Bank of Montreal Scholarship for the Next Generation of Scientists.

## Funding and additional information

This research was funded by NSERC Discovery Grants awarded to WZ (RGPIN-2019-05721) and YT (RGPIN-2017-06520). The Structural Genomics Consortium is a United Kingdom registered charity (no. 1097737) that receives funds from AbbVie, Bayer, Boehringer Ingelheim, Canada Foundation for Innovation, Eshelman Institute for Innovation, Genome Canada through the Ontario Genomics Institute (OGI-055), EU Innovative Medicines Initiative (ULTRA-DD Grant 115766), Janssen, Merck KGaA, MSD, Novartis, Ontario Ministry of Research, Innovation and Science, Pfizer, São Paulo Research Foundation-FAPESP, Takeda, and the Wellcome Trust. Y.G. is a visiting scholar sponsored by the Excellent Young Teachers program and Innovation Fund of Guangdong Ocean University, China.

## References

1. Hu, H., and Sun, S. C. (2016) Ubiquitin signaling in immune responses. Cell Res. 26, 457–483

2. Huang, T. T., and D’Andrea, A. D. (2006) Regulation of DNA repair by ubiquitylation. Nat. Rev. Mol. Cell Biol. 7, 323–334

3. Nakayama, K. I., and Nakayama, K. (2006) Ubiquitin ligases: Cell-cycle control and cancer. Nat. Rev. Cancer. 6, 369–381

4. Swatek, K. N., and Komander, D. (2016) Ubiquitin modifications. Cell Res. 26, 399–422

5. Chen, Z. J., and Sun, L. J. (2009) Nonproteolytic Functions of Ubiquitin in Cell Signaling. Mol. Cell. 33, 275–286

6. Harrigan, J. A., Jacq, X., Martin, N. M., and Jackson, S. P. (2018) Deubiquitylating enzymes and drug discovery: Emerging opportunities. Nat. Rev. Drug Discov. 17, 57–77

7. Lill, J. R., and Wertz, I. E. (2014) Toward understanding ubiquitin-modifying enzymes: From pharmacological targeting to proteomics. Trends Pharmacol. Sci. 35, 187–207

8. Abdul Rehman, S. A., Kristariyanto, Y. A., Choi, S. Y., Nkosi, P. J., Weidlich, S., Labib, K., Hofmann, K., and Kulathu, Y. (2016) MINDY-1 Is a Member of an Evolutionarily Conserved and Structurally Distinct New Family of Deubiquitinating Enzymes. Mol. Cell. 63, 146–155

9. Coleman, K. E., and Huang, T. T. (2018) In a Class of Its Own: A New Family of Deubiquitinases Promotes Genome Stability. Mol. Cell. 70, 1–3

10. Kemp, M. (2016) Recent Advances in the Discovery of Deubiquitinating Enzyme Inhibitors. Prog. Med. Chem. 55, 149–192

11. Arrowsmith, C. H., Audia, J. E., Austin, C., Baell, J., Bennett, J., Blagg, J., Bountra, C., Brennan, P. E., Brown, P. J., Bunnage, M. E., Buser-Doepner, C., Campbell, R. M., Carter, A. J., Cohen, P., Copeland, R. A., Cravatt, B., Dahlin, J. L., Dhanak, D., Edwards, A. M., Frye, S. V., Gray, N., Grimshaw, C. E., Hepworth, D., Howe, T., Huber, K. V. M., Jin, J., Knapp, S., Kotz, J. D., Kruger, R. G., Lowe, D., Mader, M. M., Marsden, B., Mueller-Fahrnow, A., Müller, S., O’Hagan, R. C., Overington, J. P., Owen, D. R., Rosenberg, S. H., Roth, B., Ross, R., Schapira, M., Schreiber, S. L., Shoichet, B., Sundström, M., Superti-Furga, G., Taunton, J., Toledo-Sherman, L., Walpole, C., Walters, M. A., Willson, T. M., Workman, P., Young, R. N., and Zuercher, W. J. (2015) The promise and peril of chemical probes. Nat. Chem. Biol. 11, 536–541

12. Zhang, W., and Sidhu, S. S. (2018) Drug development: Allosteric inhibitors hit USP7 hard. Nat. Chem. Biol. 14, 110–111

13. Colland, F., Formstecher, E., Jacq, X., Reverdy, C., Planquette, C., Conrath, S., Trouplin, V., Bianchi, J., Aushev, V. N., Camonis, J., Calabrese, A., Borg-Capra, C., Sippl, W., Collura, V., Boissy, G., Rain, J. C., Guedat, P., Delansorne, R., and Daviet, L. (2009) Small-molecule inhibitor of USP7/HAUSP ubiquitin protease stabilizes and activates p53 in cells. Mol. Cancer Ther. 8, 2286–2295

14. Gavory, G., O’Dowd, C. R., Helm, M. D., Flasz, J., Arkoudis, E., Dossang, A., Hughes, C., Cassidy, E., McClelland, K., Odrzywol, E., Page, N., Barker, O., Miel, H., and Harrison, T. (2018) Discovery and characterization of highly potent and selective allosteric USP7 inhibitors. Nat. Chem. Biol. 14, 118–125

15. Turnbull, A. P., Ioannidis, S., Krajewski, W. W., Pinto-Fernandez, A., Heride, C., Martin, A. C. L., Tonkin, L. M., Townsend, E. C., Buker, S. M., Lancia, D. R., Caravella, J. A., Toms, A. V., Charlton, T. M., Lahdenranta, J., Wilker, E., Follows, B. C., Evans, N. J., Stead, L., Alli, C., Zarayskiy, V. V., Talbot, A. C., Buckmelter, A. J., Wang, M., McKinnon, C. L., Saab, F., McGouran, J. F., Century, H., Gersch, M., Pittman, M. S., Gary Marshall, C., Raynham, T. M., Simcox, M., Stewart, L. M. D., McLoughlin, S. B., Escobedo, J. A., Bair, K. W., Dinsmore, C. J., Hammonds, T. R., Kim, S., Urbé, S., Clague, M. J., Kessler, B. M., and Komander, D. (2017) Molecular basis of USP7 inhibition by selective small-molecule inhibitors. Nature. 550, 481–486

16. Kategaya, L., Di Lello, P., Rougé, L., Pastor, R., Clark, K. R., Drummond, J., Kleinheinz, T., Lin, E., Upton, J. P., Prakash, S., Heideker, J., McCleland, M., Ritorto, M. S., Alessi, D. R., Trost, M., Bainbridge, T. W., Kwok, M. C. M., Ma, T. P., Stiffler, Z., Brasher, B., Tang, Y., Jaishankar, P., Hearn, B. R., Renslo, A. R., Arkin, M. R., Cohen, F., Yu, K., Peale, F., Gnad, F., Chang, M. T., Klijn, C., Blackwood, E., Martin, S. E., Forrest, W. F., Ernst, J. A., Ndubaku, C., Wang, X., Beresini, M. H., Tsui, V., Schwerdtfeger, C., Blake, R. A., Murray, J., Maurer, T., and Wertz, I. E. (2017) USP7 small-molecule inhibitors interfere with ubiquitin binding, Nature Publishing Group, 10.1038/nature24006

17. Clague, M. J., and Urbé, S. (2006) Endocytosis: the DUB version. Trends Cell Biol. 16, 551–559

18. Shrestha, R. K., Ronau, J. A., Davies, C. W., Guenette, R. G., Strieter, E. R., Paul, L. N., and Das, C. (2014) Insights into the mechanism of deubiquitination by jamm deubiquitinases from cocrystal structures of the enzyme with the substrate and product. Biochemistry. 53, 3199–3217

19. Nijman, S. M. B., Luna-Vargas, M. P. A., Velds, A., Brummelkamp, T. R., Dirac, A. M. G., Sixma, T. K., and Bernards, R. (2005) A genomic and functional inventory of deubiquitinating enzymes. Cell. 123, 773–786

20. Zhu, P., Zhou, W., Wang, J., Puc, J., Ohgi, K. A., Erdjument-Bromage, H., Tempst, P., Glass, C. K., and Rosenfeld, M. G. (2007) A Histone H2A Deubiquitinase Complex Coordinating Histone Acetylation and H1 Dissociation in Transcriptional Regulation. Mol. Cell. 27, 609–621

21. Zeqiraj, E., Tian, L., Piggott, C. A., Pillon, M. C., Duffy, N. M., Ceccarelli, D. F., Keszei, A. F. A., Lorenzen, K., Kurinov, I., Orlicky, S., Gish, G. D., Heck, A. J. R., Guarné, A., Greenberg, R. A., and Sicheri, F. (2015) Higher-Order Assembly of BRCC36-KIAA0157 Is Required for DUB Activity and Biological Function. Mol. Cell. 59, 970–983

22. Kweon, S. M., Chen, Y., Moon, E., Kvederaviciutė, K., Klimasauskas, S., and Feldman, D. E. (2019) An Adversarial DNA N6-Methyladenine-Sensor Network Preserves Polycomb Silencing. Mol. Cell. 74, 1138–1147.e6

23. Cooper, E. M., Cutcliffe, C., Kristiansen, T. Z., Pandey, A., Pickart, C. M., and Cohen, R. E. (2009) K63-specific deubiquitination by two JAMM/MPN+ complexes: BRISC-associated Brcc36 and proteasomal Poh1. EMBO J. 28, 621–631

24. Lauinger, L., Li, J., Shostak, A., Cemel, I. A., Ha, N., Zhang, Y., Merkl, P. E., Obermeyer, S., Stankovic-Valentin, N., Schafmeier, T., Wever, W. J., Bowers, A. A., Carter, K. P., Palmer, A. E., Tschochner, H., Melchior, F., Deshaies, R. J., Brunner, M., and Diernfellner, A. (2017) Thiolutin is a zinc chelator that inhibits the Rpn11 and other JAMM metalloproteases. Nat. Chem. Biol. 13, 709–714

25. Li, J., Yakushi, T., Parlati, F., MacKinnon, A. L., Perez, C., Ma, Y., Carter, K. P., Colayco, S., Magnuson, G., Brown, B., Nguyen, K., Vasile, S., Suyama, E., Smith, L. H., Sergienko, E., Pinkerton, A. B., Chung, T. D. Y., Palmer, A. E., Pass, I., Hess, S., Cohen, S. M., and Deshaies, R. J. (2017) Capzimin is a potent and specific inhibitor of proteasome isopeptidase Rpn11. Nat. Chem. Biol. 13, 486–493

26. Schlierf, A., Altmann, E., Quancard, J., Jefferson, A. B., Assenberg, R., Renatus, M., Jones, M., Hassiepen, U., Schaefer, M., Kiffe, M., Weiss, A., Wiesmann, C., Sedrani, R., Eder, J., and Martoglio, B. (2016) Targeted inhibition of the COP9 signalosome for treatment of cancer. Nat. Commun. 10.1038/ncomms13166

27. Raiborg, C., and Stenmark, H. (2009) The ESCRT machinery in endosomal sorting of ubiquitylated membrane proteins. Nature. 458, 445–452

28. McDonell, L. M., Mirzaa, G. M., Alcantara, D., Schwartzentruber, J., Carter, M. T., Lee, L. J., Clericuzio, C. L., Graham, J. M., Morris-Rosendahl, D. J., Polster, T., Acsadi, G., Townshend, S., Williams, S., Halbert, A., Isidor, B., David, A., Smyser, C. D., Paciorkowski, A. R., Willing, M., Woulfe, J., Das, S., Beaulieu, C. L., Marcadier, J., Geraghty, M. T., Frey, B. J., Majewski, J., Bulman, D. E., Dobyns, W. B., O’Driscoll, M., and Boycott, K. M. (2013) Mutations in STAMBP, encoding a deubiquitinating enzyme, cause microcephaly-capillary malformation syndrome. Nat. Genet. 45, 556–562

29. Kim, M. S., Kim, J. A., Song, H. K., and Jeon, H. (2006) STAM-AMSH interaction facilitates the deubiquitination activity in the C-terminal AMSH. Biochem. Biophys. Res. Commun. 351, 612–618

30. Kikuchi, K., Ishii, N., Asao, H., and Sugamura, K. (2003) Identification of AMSH-LP containing a Jab1/MPN domain metalloenzyme motif. Biochem. Biophys. Res. Commun. 306, 637–643

31. Nakamura, M., Tanaka, N., Kitamura, N., and Komada, M. (2006) Clathrin anchors deubiquitinating enzymes, AMSH and AMSH-like protein, on early endosomes. Genes to Cells. 11, 593–606

32. Sato, Y., Yoshikawa, A., Yamagata, A., Mimura, H., Yamashita, M., Ookata, K., Nureki, O., Iwai, K., Komada, M., and Fukai, S. (2008) Structural basis for specific cleavage of Lys 63-linked polyubiquitin chains. Nature. 455, 358–362

33. Bednash, J. S., Weathington, N., Londino, J., Rojas, M., Gulick, D. L., Fort, R., Han, S. H., McKelvey, A. C., Chen, B. B., and Mallampalli, R. K. (2017) Targeting the deubiquitinase STAMBP inhibits NALP7 inflammasome activity. Nat. Commun. 10.1038/ncomms15203

34. Ernst, A., Avvakumov, G., Tong, J., Fan, Y., Zhao, Y., Alberts, P., Persaud, A., Walker, J. R., Neculai, A. M., Neculai, D., Vorobyov, A., Garg, P., Beatty, L., Chan, P. K., Juang, Y. C., Landry, M. C., Yeh, C., Zeqiraj, E., Karamboulas, K., Allali-Hassani, A., Vedadi, M., Tyers, M., Moffat, J., Sicheri, F., Pelletier, L., Durocher, D., Raught, B., Rotin, D., Yang, J., Moran, M. F., Dhe-Paganon, S., and Sidhu, S. S. (2013) A strategy for modulation of enzymes in the ubiquitin system. Science (80-. ). 339, 590–595

35. Komander, D., and Rape, M. (2012) The ubiquitin code. Annu. Rev. Biochem. 81, 203–229

36. Zhang, W., Sartori, M. A., Makhnevych, T., Federowicz, K. E., Dong, X., Liu, L., Nim, S., Dong, A., Yang, J., Li, Y., Haddad, D., Ernst, A., Heerding, D., Tong, Y., Moffat, J., and Sidhu, S. S. (2017) Generation and Validation of Intracellular Ubiquitin Variant Inhibitors for USP7 and USP10. J. Mol. Biol. 429, 3546–3560

37. Krissinel, E., and Henrick, K. (2007) Inference of Macromolecular Assemblies from Crystalline State. J. Mol. Biol. 372, 774–797

38. Baiady, N., Padala, P., Mashahreh, B., Cohen-Kfir, E., Todd, E. A., Du Pont, K. E., Berndsen, C. E., and Wiener, R. (2016) The Vps27/Hrs/STAM (VHS) domain of the signaltransducing adaptor molecule (STAM) directs associated molecule with the SH3 domain of STAM (AMSH) specificity to longer ubiquitin chains and dictates the position of cleavage. J. Biol. Chem. 291, 2033–2042

39. Hologne, M., Cantrelle, F. X., Riviere, G., Guillière, F., Trivelli, X., and Walker, O. (2016) NMR Reveals the Interplay among the AMSH SH3 Binding Motif, STAM2, and Lys63- Linked Diubiquitin. J. Mol. Biol. 428, 4544–4558

40. Davies, C. W., Paul, L. N., and Das, C. (2013) Mechanism of recruitment and activation of the endosome-associated deubiquitinase AMSH. Biochemistry. 52, 7818–7829

41. Ernst, A., Avvakumov, G., Tong, J., Fan, Y., Zhao, Y., Alberts, P., Persaud, A., Walker, J. R., Neculai, A. M., Neculai, D., Vorobyov, A., Garg, P., Beatty, L., Chan, P. K., Juang, Y. C., Landry, M. C., Yeh, C., Zeqiraj, E., Karamboulas, K., Allali-Hassani, A., Vedadi, M., Tyers, M., Moffat, J., Sicheri, F., Pelletier, L., Durocher, D., Raught, B., Rotin, D., Yang, J., Moran, M. F., Dhe-Paganon, S., and Sidhu, S. S. (2013) Supplementary Material for - A strategy for modulation of enzymes in the ubiquitin system. Science (80-. ). 339, 590–595

42. Garg, P., Ceccarelli, D. F., Keszei, A. F. A., Kurinov, I., Sicheri, F., and Sidhu, S. S. (2020) Structural and Functional Analysis of Ubiquitin-based Inhibitors That Target the Backsides of E2 Enzymes. J. Mol. Biol. 432, 952–966

43. Gabrielsen, M., Buetow, L., Nakasone, M. A., Ahmed, S. F., Sibbet, G. J., Smith, B. O., Zhang, W., Sidhu, S. S., and Huang, D. T. (2017) A General Strategy for Discovery of Inhibitors and Activators of RING and U-box E3 Ligases with Ubiquitin Variants. Mol. Cell. 68, 456–470.e10

44. Zhang, W., Wu, K. P., Sartori, M. A., Kamadurai, H. B., Ordureau, A., Jiang, C., Mercredi, P. Y., Murchie, R., Hu, J., Persaud, A., Mukherjee, M., Li, N., Doye, A., Walker, J. R., Sheng, Y., Hao, Z., Li, Y., Brown, K. R., Lemichez, E., Chen, J., Tong, Y., Harper, J. W., Moffat, J., Rotin, D., Schulman, B. A., and Sidhu, S. S. (2016) System-Wide Modulation of HECT E3 Ligases with Selective Ubiquitin Variant Probes. Mol. Cell. 62, 121–136

45. Gorelik, M., Orlicky, S., Sartori, M. A., Tang, X., Marcon, E., Kurinov, I., Greenblatt, J. F., Tyers, M., Moffat, J., Sicheri, F., and Sidhu, S. S. (2016) Inhibition of SCF ubiquitin ligases by engineered ubiquitin variants that target the Cul1 binding site on the Skp1-F-box interface. Proc. Natl. Acad. Sci. U. S. A. 113, 3527–3532

46. Manczyk, N., Veggiani, G., Teyra, J., Strilchuk, A. W., Sidhu, S. S., and Sicheri, F. (2019) The ubiquitin interacting motifs of USP37 act on the proximal Ub of a di-Ub chain to enhance catalytic efficiency. Sci. Rep. 9, 1–11

47. Teyra, J., Singer, A. U., Schmitges, F. W., Jaynes, P., Kit Leng Lui, S., Polyak, M. J., Fodil, N., Krieger, J. R., Tong, J., Schwerdtfeger, C., Brasher, B. B., Ceccarelli, D. F. J., Moffat, J., Sicheri, F., Moran, M. F., Gros, P., Eichhorn, P. J. A., Lenter, M., Boehmelt, G., and Sidhu, S. S. (2019) Structural and Functional Characterization of Ubiquitin Variant Inhibitors of USP15. Structure. 27, 590–605.e5

48. Zhang, W., Bailey-Elkin, B. A., Knaap, R. C. M., Khare, B., Dalebout, T. J., Johnson, G. G., van Kasteren, P. B., McLeish, N. J., Gu, J., He, W., Kikkert, M., Mark, B. L., and Sidhu, S. S. (2017) Potent and selective inhibition of pathogenic viruses by engineered ubiquitin variants. PLOS Pathog. 13, e1006372

49. Mund, T., Lewis, M. J., Maslen, S., and Pelham, H. R. (2014) Peptide and small molecule inhibitors of HECT-type ubiquitin ligases. Proc. Natl. Acad. Sci. U. S. A. 111, 16736–16741

50. Gorelik, M., Manczyk, N., Pavlenco, A., Kurinov, I., Sidhu, S. S., and Sicheri, F. (2018) A Structure-Based Strategy for Engineering Selective Ubiquitin Variant Inhibitors of Skp1-Cul1-F-Box Ubiquitin Ligases. Structure. 26, 1226–1236.e3

51. Dikic, I., Wakatsuki, S., and Walters, K. J. (2009) Ubiquitin-binding domains from structures to functions. Nat. Rev. Mol. Cell Biol. 10, 659–671

52. Manczyk, N., Yates, B. P., Veggiani, G., Ernst, A., Sicheri, F., and Sidhu, S. S. (2017) Structural and functional characterization of a ubiquitin variant engineered for tight and specific binding to an alpha-helical ubiquitin interacting motif. Protein Sci. 26, 1060–1069

53. Manczyk, N., Veggiani, G., Gish, G. D., Yates, B. P., Ernst, A., Sidhu, S. S., and Sicheri, F. (2019) Dimerization of a ubiquitin variant leads to high affinity interactions with a ubiquitin interacting motif. Protein Sci. 28, 848–856

54. Walden, M., Tian, L., Ross, R. L., Sykora, U. M., Byrne, D. P., Hesketh, E. L., Masandi, S. K., Cassel, J., George, R., Ault, J. R., El Oualid, F., Pawłowski, K., Salvino, J. M., Eyers, P. A., Ranson, N. A., Del Galdo, F., Greenberg, R. A., and Zeqiraj, E. (2019) Metabolic control of BRISC–SHMT2 assembly regulates immune signalling. Nature. 570, 194–199

55. Paudel, P., Zhang, Q., Leung, C., Greenberg, H. C., Guo, Y., Chern, Y. H., Dong, A., Li, Y., Vedadi, M., Zhuang, Z., and Tong, Y. (2019) Crystal structure and activity-based labeling reveal the mechanisms for linkage-specific substrate recognition by deubiquitinase USP9X. Proc. Natl. Acad. Sci. U. S. A. 116, 7288–7297

56. Grochulski, P., Fodje, M. N., Gorin, J., Labiuk, S. L., and Berg, R. (2011) Beamline 08ID-1, the prime beamline of the Canadian macromolecular crystallography facility. J. Synchrotron Radiat. 18, 681–684

57. Minor, W., Cymborowski, M., Otwinowski, Z., and Chruszcz, M. (2006) HKL-3000: The integration of data reduction and structure solution - From diffraction images to an initial model in minutes. Acta Crystallogr. Sect. D Biol. Crystallogr. 62, 859–866

58. Tonikian, R., Zhang, Y., Boone, C., and Sidhu, S. S. (2007) Identifying specificity profiles for peptide recognition modules from phage-displayed peptide libraries. Nat. Protoc. 2, 1368–1386

